# Self-esteem modulates beneficial causal attributions in the formation of novel self-beliefs

**DOI:** 10.1101/2025.10.27.684784

**Authors:** Annalina V. Mayer, Alexander Schröder, David S Stolz, Nora Czekalla, Frieder M Paulus, Laura Müller-Pinzler, Sören Krach, Tobias Kube

## Abstract

Healthy individuals typically attribute successes to internal causes, such as their abilities, and failures to external factors, like bad luck. In contrast, individuals with depression and low self-esteem are more likely to attribute failures to internal causes and successes to external causes. At the same time, depression and low self-esteem are associated with negatively biased self-related learning and self-beliefs. Although causal attributions have been shown to influence belief formation and updating, the dynamic interaction between real-time attributions and self-related learning remains poorly understood. In this study, we used a validated self-related learning task to investigate how internal versus external attributions of performance feedback affect the formation of self-beliefs and how these processes relate to depressive symptoms and self-esteem. Drawing on a computational model that incorporates prediction error valence and causal attributions, we found that participants updated their self-beliefs less when feedback was attributed to external causes. Furthermore, individuals with higher levels of depression and lower self-esteem showed a stronger negativity bias in learning. Lower self-esteem was also linked to a reduced self-serving bias in attributions. These findings provide insight into the cognitive mechanisms that may contribute to the development and maintenance of negative self-beliefs commonly observed in depression.

## Introduction

Attributional styles, referred to as individual tendencies to explain the causes of events, play a crucial role in shaping people’s beliefs and their understanding of themselves and the world. In particular, attributional styles reflect whether individuals interpret life events as resulting from internal or external, stable or unstable, and global or specific causes (Abramson et al., 1978). Resembling recent computational considerations on links between belief formation and attributions (Zamfir & Dayan, 2022), a person with a pessimistic attributional style may view negative events as internally caused, enduring, and pervasive. Conversely, an optimistic attributional style tends to foster resilience by attributing setbacks to external, temporary, and situation-specific factors (Sweeney et al., 1986). Understanding attributional styles thus offers critical insight into cognitive vulnerability and resilience across various aspects of psychological functioning.

Attributional styles have been closely linked to mental health outcomes (Cheng & Furnham, 2001). For instance, while healthy individuals tend to attribute successes to internal causes (e.g., their abilities) and failures to external causes (e.g., bad luck) (Larson, 1977), individuals with depression often show the opposite pattern (Kuiper, 1978; Peterson & Seligman, 1984; Sweeney et al., 1986). In fact, a pessimistic attributional style is associated with low self-esteem (Fitch, 1970; Romney, 1994; Tennen et al., 1987) and is considered a key cognitive factor in the development and maintenance of depression and other mental disorders (Abramson et al., 1978; Fresco et al., 2006; Heimberg et al., 1989; Huang, 2015; Southall & Roberts, 2002).

Traditionally, attributional styles have been studied as interindividual traits, with much of the research examining the effects of causal attributions on emotion, motivation, and mental health relying on self-report questionnaires (Elig & Frieze, 1979). However, there is limited understanding of how real-time causal attributions of successes and failures interact with self-related learning, although recent computational considerations highlight the importance of this particular interplay (Bentall et al., 2001; Zamfir & Dayan, 2022). This is particularly relevant for understanding the cognitive mechanisms that underlie the negatively biased self-beliefs characteristic of depression (Beck, 1979).

In this study, we aim to answer the following questions: First, do momentary attributions of failures and successes generally influence self-related learning, and thus the formation of self-related beliefs? Second, are depressive symptoms and self-esteem related to different attributional patterns during self-related learning, and does this contribute to biased self-beliefs?

A growing body of research has demonstrated that depression and low self-esteem are associated with negatively biased self-related learning. For example, individuals with major depressive disorder show difficulties in updating their negative beliefs after unexpected positive experiences (Czekalla et al., 2024; Kube et al., 2019, 2020). Individuals with clinical depression and those with elevated depression levels were also found to interpret ambiguous situations more negatively and less positively, especially when they contain self-relevant information (Everaert et al., 2017). Further, while healthy individuals tend to update their expectations about future events in an optimistically biased way, individuals with depression do not (Garrett et al., 2014; Korn et al., 2014). This absence of an optimistic update bias has recently also been shown during learning from social evaluation, where higher levels of depressive symptoms were associated with a blunted response to positive social feedback (Hoffmann et al., 2024). Similarly, individuals with low self-esteem tend to learn less from social feedback than those with high self-esteem and are more likely to expect others to dislike them (Will et al., 2020). Low self-esteem is also linked to a stronger negativity bias when learning about one’s own abilities (Müller-Pinzler et al., 2022, 2019).

Biased self-beliefs in depression and individuals with low self-esteem might be related to differences in attributional styles, since causal attributions may influence how beliefs are formed and updated. To illustrate this, consider the following example: A good student receives a poor grade on a math exam. How the student interprets this outcome, whether as a result of internal or external causes, shapes how they think about their mathematical ability. If the student believes the low grade was due to their lack of understanding of the material (internal attribution), they are more likely to lower their belief about their ability. On the other hand, if the student attributes the poor grade to the exam being unfair or the teacher being biased (external attribution), they may dismiss the feedback and continue to hold a high belief about their ability. If the student has the tendency to attribute failures to internal causes, these failures are more likely to shape their self-beliefs than successes, resulting in negatively biased beliefs.

A number of behavioral studies have demonstrated this relationship of causal attributions and subsequent self-beliefs in a performance context: when participants attributed a successful performance to internal causes (i.e., their ability), this heightened their self-efficacy, while attributions to external causes (i.e., good luck) lowered their self-efficacy (Silver et al., 1995; Stajkovic & Sommer, 2000). More recent advancements in neuroscience and psychology have applied computational modeling techniques to explore how causal attributions shape beliefs about action-outcome relationships in a reinforcement-learning task (Dorfman et al., 2019, 2021). Participants gave less weight to outcomes they believed were caused by external factors – the interference of a hidden agent – rather than their own actions. In other words, when participants thought an outcome was manipulated, they learned less from it. This process resulted in biased learning depending on whether the hidden agent changed the outcome positively or negatively (Dorfman et al., 2019, 2021).

We aimed to expand this research by examining the role of causal attributions during the formation of self-beliefs (Krach et al., 2024) rather than beliefs about the world (Dorfman et al., 2019, 2021). Using a validated learning task (Czekalla et al., 2021; Müller-Pinzler et al., 2022, 2019; Schröder et al., 2024), we investigated how internal vs. external attributions of performance feedback shape self-related learning in novel ability domains. Drawing on validated computational learning models that account for prediction error valence (Czekalla et al., 2021; Müller-Pinzler et al., 2022, 2019; Schröder et al., 2024) and that have been further extended to consider causal attributions, we first hypothesized that participants would learn less from feedback attributed to external causes compared to feedback attributed to their own abilities. Second, we hypothesized that depressive symptom severity and self-esteem would be associated with biases in learning and attributions. Specifically, based on research pointing to deficits in learning from novel positive experiences in people with elevated levels of depression (Korn et al., 2014; Kube & Eggers, 2025; Kube et al., 2019), we expected that participants with higher levels of depression would show a more negative bias in how they learned from the feedback. Similarly, we expected that lower self-esteem would be linked to a more negative bias, as shown before (Müller-Pinzler et al., 2022, 2019). We also anticipated that with increasing depressive symptom severity individuals would more likely attribute failures, defined as worse-than-expected feedback, to internal causes, while attributing successes, defined as better-than-expected feedback, to external causes, as implicated by the attributional theory of depression (Abramson et al., 1978; Fresco et al., 2006; Heimberg et al., 1989; Huang, 2015; Southall & Roberts, 2002).

## Results

### Measuring self-belief formation and attributions of self-related feedback

Sixty-four university students (75.8% female) aged between 19 and 33 years (M = 22.8, SD = 3.3) completed an adapted version of a validated self-related learning task, the Learning Of Own Performance (LOOP) task (Müller-Pinzler et al., 2022, 2019). In brief, the LOOP task allows participants to incrementally form beliefs about their ability to estimate various properties, such as the height of houses or the weight of animals. This is done by presenting predefined feedback after participants perform an estimation task (Figure 1A). Before each estimation question, participants are asked to rate their expected performance. This is done to assess whether participants adapt their expectations to the received feedback, thus capturing the belief formation process (see Methods for details). In this study, we included a condition during which participants could attribute their performance feedback to an external “agent” (computer) rather than their own ability (Figure 1B). This was implemented by informing participants that, in some trials, their feedback might be manipulated by the computer (see Figure 1A). In these trials, participants were asked to indicate whether they thought the current feedback had been manipulated, which allowed us to assess whether the feedback was internally or externally attributed. Participants were not informed beforehand about the frequency or degree of the alleged manipulation, in line with Dorfman et al. (2019). After the experiment, depressive symptoms were measured using the Patient Health Questionnaire (PHQ-9) (Kocalevent et al., 2013; Kroenke et al., 2001). We also assessed self-esteem using the Self-Description Questionnaire (SDQ-III) (Marsh & O’Neill, 1984).

**Figure 1:**
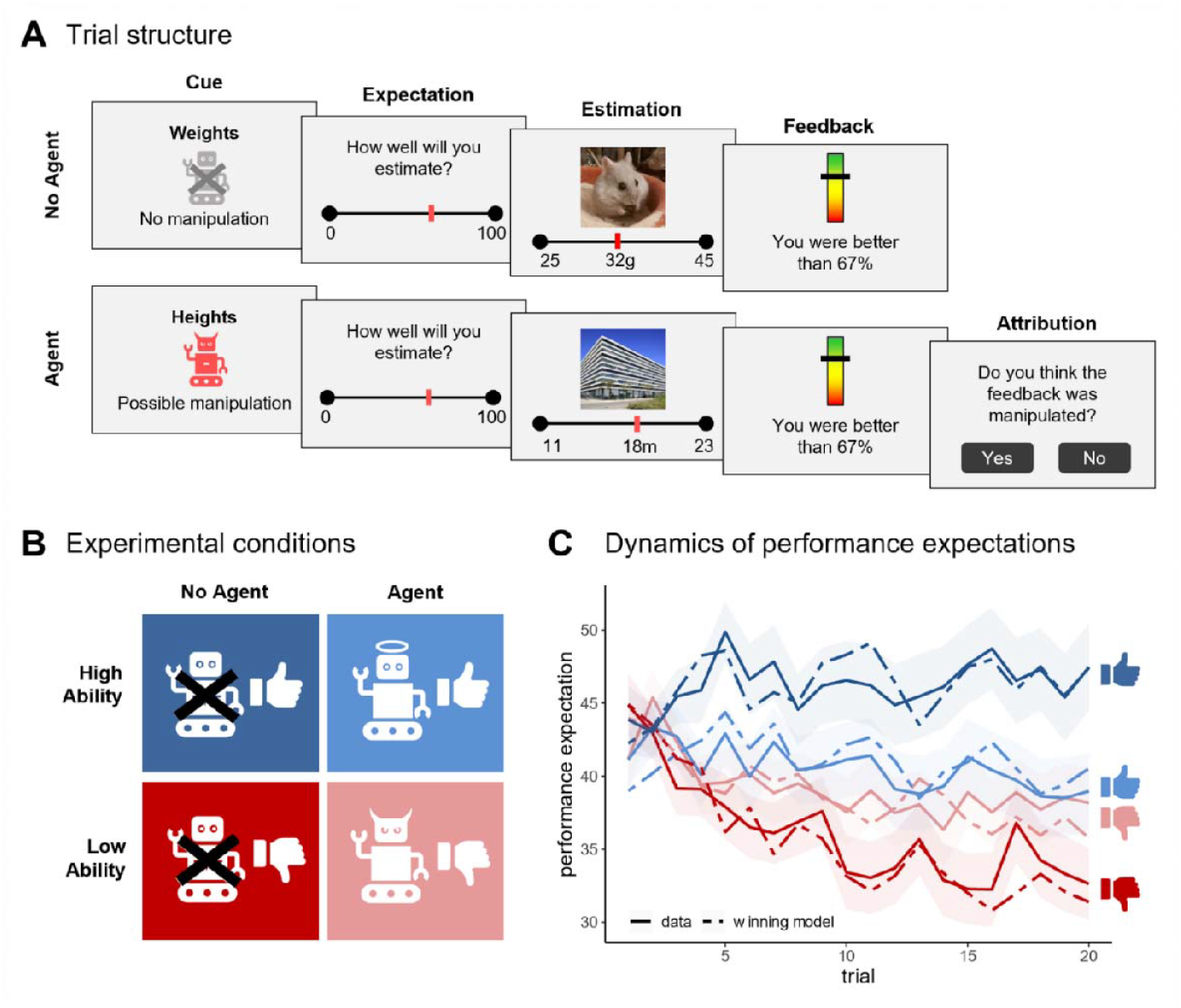
Overview of the Learning Of Own Performance (LOOP) task and task performance. **A:** In a within-subject design, participants were asked to estimate different properties in four distinct categories. A cue indicating the estimation category and whether a manipulation of the feedback was possible was followed by a performance expectation rating. The estimation question was subsequently presented for 10⍰s. Feedback was presented for 5 s after each estimation, indicating the participant’s performance compared to an alleged reference group. In trials of the Agent condition, participants were asked whether they believed the feedback had been manipulated. **B:** The four estimation categories were randomly assigned to four experimental conditions. Two categories were randomly paired with mostly positive feedback (High Ability) and two with mostly negative feedback (Low Ability). Participants were informed that in two categories, their feedback could be manipulated by the computer (Agent), such that feedback in one category might be improved and in the other worsened. In the other two categories, the feedback was allegedly not manipulated (No Agent). **C:** Participants adjusted their performance expectations (solid lines) based on the feedback they received, with the adjustments varying depending on whether a manipulation was possible. Shaded areas represent the standard errors for the actual performance expectations for each trial. Our winning computational model captured the dynamics in participants’ performance expectations (dashed line).

### Model-agnostic behavioral analysis

First, we explored whether participants’ performance expectations varied depending on the feedback condition (Ability: High vs. Low) and possible manipulation by the agent (Interference: Agent vs. No Agent) over the course of the experiment. Results of a repeated-measures ANOVA yielded significant main effects of Trial (*F*(1,63) = 6.30, *p* =.015, η^2^_G_ = 0.01) and Ability (*F*(1,63) = 18.83, *p* <.001, η^2^_G_ = 0.08), as well as a significant Trial × Ability interaction (*F*(1,63) = 17.74, *p* <.001, η^2^_G_ = 0.01), indicating that participants adjusted their expectations over time depending on the provided feedback. Importantly, a significant two-way Ability × Interference interaction (*F*(1,63) = 10.15, *p* =.002, η^2^_G_ = 0.05) as well as a significant three-way Trial × Ability × Interference interaction (*F*(1,63) = 16.17, *p* <.001, η^2^_G_ = 0.01) suggested that changes in performance expectations were not only influenced by the feedback condition, but also by whether a manipulation by the agent was possible (Figure 1C).

### Selection of computational models of self-belief formation

Participants’ trial-by-trial updating of performance expectations was quantified using computational modeling. Changes in performance expectations were modeled through updates from prediction errors (PEs) using delta-rule reinforcement learning models based on Rescorla and Wagner (1972). The model that best described the observed behavior contained separate learning rates (α) for updates following positive vs. negative PEs (see Figure 2B and Methods for model space and equations of the winning model). This allowed a PE-valence-based description of belief formation. The winning model also included a weighting factor, *w*, which reduced the updates in response to extreme feedback (towards 0% or 100%), assuming that those extremes are considered less probable and therefore less informative (Czekalla et al., 2024; Müller-Pinzler et al., 2022; Schröder et al., 2024). Further, the winning model included an attribution weight factor, *s*, that scales how strongly participants learn from externally attributed feedback, relative to the rate at which they learn from internally attributed feedback. An *s* factor closer to 0 indicates that on trials where subjects attribute feedback externally they learn at a similar rate as on trials where feedback is attributed internally. Inversely, an *s* factor of (close to) 1 indicates that subjects do not learn from externally attributed feedback at all, thus maintaining their current self-belief (see Supplementary Figure S2 for an illustration of this effect).

**Figure 2:**
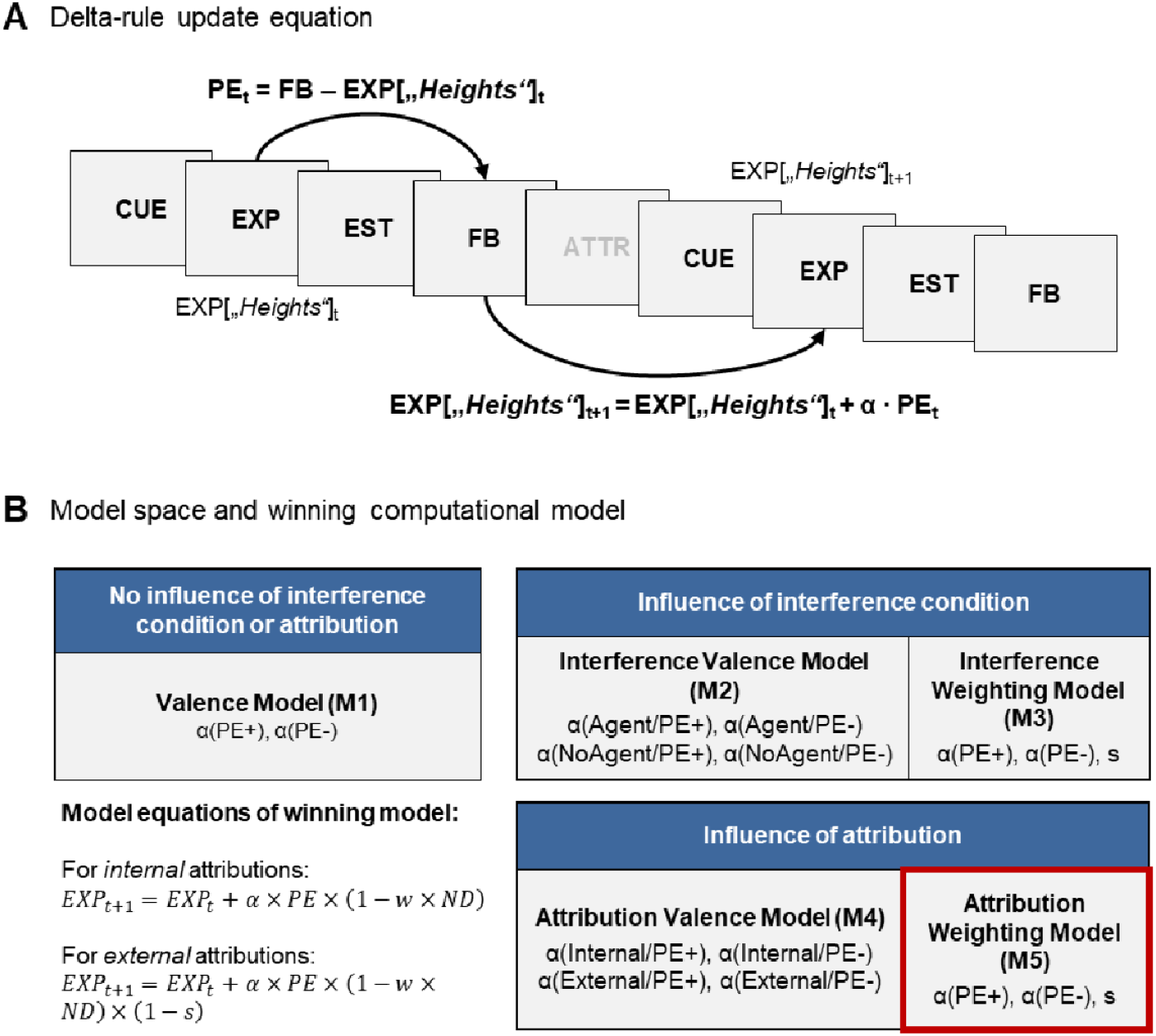
Computational modeling of self-belief formation. **A:** All computational models in our model space were built on Rescorla-Wagner delta-rule update equations, which modeled changes in performance expectations (EXP) by incorporating multiple learning rates (α, see Methods) and accounting for trial-by-trial prediction errors (PE) based on the provided feedback (FB). **B:** In our models, we distinguished two factors that might influence learning rates and/or expectation updates in general: The Interference condition (Agent vs. No Agent) and trial-by-trial attributions (external vs. internal). As a baseline, we included a model that did not account for any of these factors (M1), but included separate learning rates for positive and negative PEs. While valence models (M2 and M4) incorporated separate valence-specific learning rates for each level of the factor of interest, weighting models (M3 and M5) included two valence-specific learning rates and an additional weight factor *s* that reduced the update depending on the attribution or interference condition (see Methods for details). The winning model is outlined in red.

Posterior predictive checks validated that the winning model accurately captured the key effects from our model-agnostic analysis, showing that the feedback and interference conditions jointly influenced dynamics in predicted performance expectations (see Supplementary Table S2 and Figure 1C).

### Negativity bias in self-belief formation and reduced updating after externally attributed feedback

On average, participants showed higher learning rates for negative (*Mdn* =.024) than for positive PEs (*Mdn* = 0.11) (Wilcoxon signed rank test, *z*(63) = -3.74, *p* <.001, *r* =.468), suggesting a general negativity bias during self-belief formation over all experimental conditions (Figure 3A). Attribution weight factors were significantly greater than 0 with a mean value of *M* = 0.51 (*SD* = 0.17, *t*(63) = 24.3, *p* <.001, *d* = 3.04, Figure 3B). Thus, on average, updates in performance expectations were reduced by half after externally attributed feedback compared to internally attributed feedback. In other words, participants integrated feedback into their self-beliefs to a lesser extent when they believed the computer had manipulated it.

**Figure 3:**
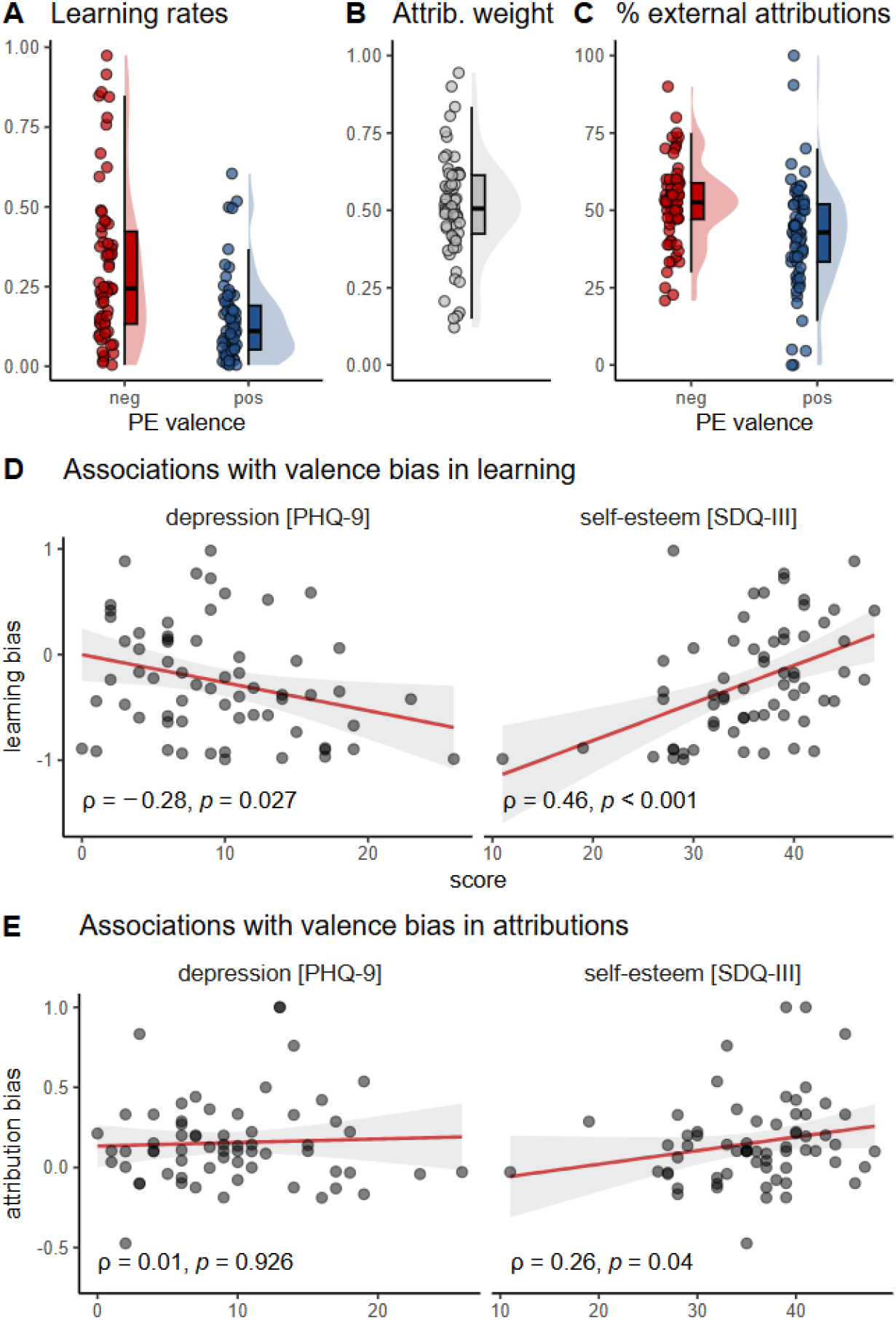
Parameters of winning model, percentage of external attributions made during the task, and associations of biases in learning and attributions with depression and self-esteem. **A:** Learning rates indicate stronger updates in performance expectations after negative vs. positive prediction errors. **B:** Attribution weight factors were significantly greater than zero, leading to diminished updates after external vs. internal attributions. **C:** The percentage of external attributions was higher after negative vs. positive prediction errors. **D:** Higher depression levels and lower self-esteem were associated with a stronger negativity bias in learning. E: Higher self-esteem was associated with a stronger self-serving bias in attributions.

Self-serving bias in attributions

Since learning rates differed significantly depending on PE valence, we explored whether trial-by-trial attributions of feedback in the Agent condition varied similarly in a valence-specific manner. Indeed, the percentage of external attributions was significantly higher after negative (*Mdn* = 52.6%) compared to positive PEs (*Mdn* = 42.9%) (Wilcoxon signed rank test, *z*(63) = -4.15, *p* <.001, *r* =.518, Figure 3C). Participants were thus more likely to believe that the feedback reflected their true ability (rather than a manipulation by the computer) when it was better than expected, compared to worse than expected, indicating a self-serving bias in attributions.

### Depression and self-esteem are associated with biased learning and attributions

To assess the relationship of depressive symptoms and self-esteem with biases in learning and attributions, we first calculated individual valence bias scores for each participant (see Methods). Correlation analyses showed a negative association between bias scores in learning and depressive symptoms (Spearman *ρ* = -.28, *p* =.027) as well as a positive association between bias scores in learning and self-esteem (*ρ* =.46, *p* <.001, Figure 3D). This suggests that individuals with elevated depressive symptoms and lower self-esteem showed a stronger negativity bias in learning. Similarly, self-esteem was positively associated with bias scores in attributions (*ρ* =.26, *p* =.04, Figure 3E), indicating that those with higher self-esteem were more biased toward attributing the feedback externally after negative PEs compared to positive PEs, thus showing a more pronounced self-serving bias. Unexpectedly, the degree of depressive symptoms was not significantly associated with valence bias scores in attributions (*ρ* =.01, *p* =.926). In general, while depression and self-esteem were strongly negatively correlated (*ρ* = -.60, *p* <.001), there was no significant correlation between the two bias types (*ρ* = -.02, *p* =.879).

## Discussion

The present study investigated how causal attributions influence the formation of self-beliefs in real-time by introducing an external “agent” that possibly manipulates performance feedback in our learning task. We replicated a negativity bias in self-related learning in a performance context, showing that participants adjusted their self-beliefs to a stronger extent after worse-than-expected feedback, compared to better-than-expected feedback (Brotzeller & Gollwitzer, 2024; Czekalla et al., 2021; Ertac, 2011; Müller-Pinzler et al., 2022, 2019; Schröder et al., 2024; Zamfir & Dayan, 2022). This was more prominent in individuals with lower self-esteem and higher depressive symptom severity. Second, causal attributions of the feedback had an impact on self-related learning. Computational modeling captured the data most accurately when individual trial-by-trial causal attributions were considered. When feedback was attributed to the external agent, updates of self-beliefs were attenuated. That is, participants adjusted their self-beliefs to a lesser degree when they thought that the feedback had been manipulated. Third, a self-serving attributional bias was identified. Participants were more likely to attribute feedback that was worse-than-expected to external factors and better-than-expected feedback to their own abilities. Here, less self-serving attributions were linked to low self-esteem, rather than the severity of depressive symptoms. This suggests that self-esteem plays a specific role in influencing attributional tendencies.

These findings extend prior work on causal attributions and associated biases by manipulating and assessing trial-by-trial causal attributions during self-related learning. Previous studies have demonstrated that people tend to attribute positive outcomes to themselves and negative outcomes to external causes. This could be shown when learning about reward probabilities in a forced-choice task (Dorfman et al., 2019, 2021), when attributing losses in a gambling task (Morewedge, 2009), and even in unambiguous contexts with clear clues for causality (Wang et al., 2017). However, these studies mainly focused on attributions of events that reflected beliefs about the world, such as causes of gains or losses in reward learning. By examining causal attributions in learning about oneself, we show that causal attributions are not only related to past outcomes but also directly influence the dynamics of trial-by-trial self-belief formation in real-time. Importantly, our results highlight two central biases in self-belief formation: A negativity bias in learning, reflected by an overemphasis on negative prediction errors compared to positive prediction errors, and a self-serving attributional bias, in which unfavorable results are more likely attributed to external causes. Although both the negativity bias (Czekalla et al., 2021; Müller-Pinzler et al., 2022, 2019; Schröder et al., 2024) and the self-serving attributional bias (Campbell & Sedikides, 1999; Dorfman et al., 2019, 2021) have been documented independently in healthy populations, our study emphasizes their concurrent occurrence. In particular, the possibility of external interference in our task introduced uncertainty about the validity of the feedback. Under these conditions, external attributions might offer a cognitive safeguard: participants could question whether the outcome truly reflected their own performance. While overall the negativity bias in learning still emerged, the attributional pattern attenuated the integration of especially worse-than-expected feedback, and this led to stagnating performance expectations in the experimental conditions where a manipulation was allegedly possible. At the same time, these attributions may help maintain self-efficacy and self-coherence (Mezulis et al., 2004; Silver et al., 1995; Stajkovic & Sommer, 2000). In addition, while self-serving attributions predominated, belief formation was also attenuated in the interference condition with better-than-expected feedback. This further emphasizes the uncertainty under which participants operated and illustrates a possible conflict of two goals: integrating feedback that shifts self-beliefs further towards desirable self-views vs. protection against possibly erroneous updates (Duval & Silvia, 2002).

The results also showed a significant role of self-esteem. Participants with higher self-esteem expressed a lesser negativity bias in learning, as was the case in previous studies (Müller-Pinzler et al., 2022, 2019). Higher self-esteem appears to protect against an overemphasis on worse-than-expected feedback. At the same time, individuals with higher self-esteem demonstrated a stronger self-serving bias in attribution, as they were more inclined to attribute negative outcomes to external factors. For these individuals, this tendency resulted in diminished updates to their self-beliefs after worse-than-expected feedback, which could have otherwise shifted their beliefs in unfavorable directions. From this perspective, again, self-serving attributions appear ambivalent: they constrain learning but serve an adaptive function by stabilizing desirable self-belief systems (Mezulis et al., 2004; Shepperd et al., 2008; Silver et al., 1995; Stajkovic & Sommer, 2000). These findings align with social-cognitive theories (Bandura, 1986, 1997; Marsh, 1990; Silver et al., 1995; Stajkovic & Sommer, 2000), which propose reciprocal reinforcement between attributional styles and self-beliefs. Such recursive dynamics may not only explain the persistence of self-serving views but also the persistence of pessimistic self-views: external attributions of positive outcomes impede corrective learning, and additionally, negative self-beliefs increase the likelihood of maladaptive, pessimistic attributions that, in the long run, may impair self-esteem (Peterson & Seligman, 1984; Sweeney et al., 1986).

Notably, self-serving motives – those to preserve self-efficacy, self-coherence, and positive affect – are also discussed in the context of the optimism bias in belief updating that was often shown in other domains of self-referential learning: the tendency of updating expectations about future events in an optimistically biased way (Korn et al., 2014; Sharot & Garrett, 2016). This supports the notion that attributional processes play an important role in the formation and updating of (self-)beliefs, especially in their role to protect desirable self-beliefs.

Contrary to our hypothesis, attributional bias was not significantly associated with depressive symptoms. However, several explanations are possible. First, the predominantly subclinical symptom levels in our sample may have had restricted variance and limited power to detect such associations. Depression, as assessed in our study using the PHQ-9 questionnaire, is a multidimensional construct that encompasses affective and somatic symptoms, which are less directly linked to attributional processes (Kroenke et al., 2001). By contrast, self-esteem as a major facet of depression (Beck, 1967; Orth & Robins, 2013) may be a more specific predictor of pessimistic attributional tendencies, given its close alignment with self-referential cognition (Duval & Silvia, 2002; Fitch, 1970; Romney, 1994; Shepperd et al., 2008; Tennen et al., 1987). Given the large body of literature that could show effects of depressive symptom severity on self-related feedback-processing (Everaert et al., 2017), belief formation and updating processes (Czekalla et al., 2024; Garrett et al., 2014; Korn et al., 2014; Kube, 2023; Kube & Eggers, 2025), and attenuated self-serving attributional bias in affected individuals (Mezulis et al., 2004), research that focuses on further disentangling involved biases and the role of causal attributions may help to provide better therapeutic interventions and, more generally, a more holistic understanding on how individuals arrive (and possibly stay) at their self-beliefs.

## Conclusion and Outlook

Our findings suggest that self-belief formation in a performance context can be shaped by two distinct, yet concurrently occurring biases. The first is a negativity bias in learning, meaning a greater emphasis on negative prediction errors compared to positive ones. The second is a self-serving attributional bias, characterized by external causal attributions of worse-than-expected feedback, which is linked to self-esteem. Individuals with lower self-esteem, who disproportionately attribute negative outcomes to their own abilities, may therefore be especially prone to developing enduring negative self-beliefs.

Future research should extend these findings to clinical samples with diagnosed major depression to further establish the relevance of attributional styles in self-belief formation for psychopathology and therapeutic interventions. More specifically, interventions that target attributional styles, for instance, by addressing pessimistic and dysfunctional attributions, may enhance corrective learning about self-related concepts and abilities and thereby reduce vulnerability to depression.

## Methods

### Preregistration and data availability

This study’s design and general research questions were preregistered before data collection (see aspredicted.org/n75w-npng.pdf). Although the general study design remained unchanged, it was necessary to make some adjustments in the testing of hypotheses, as the use of a more complex computational model was found to be more effective (see Supplementary Note 1 for more information). The collected data, results of the computational modeling approach, and the analysis scripts to reproduce the statistical results reported in this manuscript are available through the Open Science Framework (https://osf.io/2avsh/?view_only=5bd60291ceab4018b181eb26d98ca503). All variables that could be used to identify individual participants (e.g., sociodemographic data) were removed from this data set.

### Participants

The sample consisted of 66 university students (75.8% female) aged between 19 and 33 years (*M* = 22.8, *SD* = 3.3) who were recruited from the University of Kaiserslautern-Landau, Germany. All participants were fluent in German, had either normal or corrected-to-normal vision, and gave written informed consent. The study was approved by the Ethics Committee of the University of Kaiserslautern-Landau (reference number: LEK517add) and conducted in accordance with the ethical guidelines of the American Psychological Association. Two participants were excluded due to missing variance in their responses, indicating a lack of task engagement, which left a total of 64 participants to be analyzed.

### Learning Of Own Performance task

The Learning Of Own Performance (LOOP) task enables participants to incrementally learn about their alleged ability in estimating properties, for example, the height of houses or the weight of animals. The original version of the LOOP task was introduced and validated in a series of behavioral and neuroimaging studies (Czekalla et al., 2024, 2021; Müller-Pinzler et al., 2022, 2019; Schröder et al., 2024). In this study, we used an adapted version of the LOOP task including a condition that allowed participants to attribute performance feedback to an intervention by the computer (or “agent”) rather than their own ability.

On a trial-by-trial basis, participants were asked to estimate different properties and received predefined performance feedback in four distinct estimation categories (Figure 1A). Without the participant’s knowledge, two categories were randomly paired with mostly better-than-expected feedback and two with mostly worse-than-expected feedback. Participants were additionally told that in two categories, their feedback might be manipulated by the computer, such that feedback in one category might be improved and in the other worsened, without specifying the frequency or extent of this possible manipulation. In fact, participants received the same predefined feedback as in the categories where no manipulation was supposedly applied. This resulted in a 2×2 within-subject design with 20 trials per condition (Ability [High vs. Low] × Interference [Agent vs. No Agent], Figure 1B). Trials from all conditions were intermixed in a fixed order, with no more than two consecutive trials of the same condition.

Each trial started with a cue indicating the estimation category (e.g., height) and whether a manipulation of the feedback was possible (Figure 1A). Participants were then asked to state how well they expected to perform in this trial in comparison to a reference group. Following each performance expectation rating, the estimation question was presented for 10Cs. During the estimation period, continuous response scales below the pictures determined a range of plausible answers for each question. Performance feedback was provided for 5 seconds after each estimation, indicating the participant’s accuracy as percentiles compared to an alleged reference group of 350 university students, who, according to the cover story, had been tested previously (e.g., ‘You are better than 94% of the reference participants’). The feedback was determined by a series of fixed prediction errors (PEs) relative to the participant’s current belief about their abilities. This belief was calculated as the average of the last five performance expectation ratings per category, starting at 50% before participants rated their expectations. This approach resulted in varying feedback sequences across participants, while keeping PEs largely independent of individual performance expectations and ensuring a balanced distribution of negative and positive PEs across conditions. In trials of the Agent condition, participants were subsequently asked whether they believed the feedback they just received had been manipulated, thereby assessing whether they attributed the feedback to external or internal causes.

### Questionnaires and debriefing

Prior to the LOOP task, we collected general demographic information, assessed participants’ self-beliefs regarding their estimation ability, and measured self-esteem. To assess self-esteem, participants completed the general self-concept scale from the Self-Description Questionnaire-III (SDQ-III) (Marsh & O’Neill, 1984), which is based on the Rosenberg Self-Esteem Scale (Rosenberg, 1965). Depressive symptoms were assessed after the LOOP task using the Patient Health Questionnaire (PHQ-9). The PHQ-9 is the depression module of the PRIME-MD diagnostic instrument for common mental disorders and has been validated in the general German population (Gräfe et al., 2004; Kocalevent et al., 2013; Kroenke et al., 2001). After completing the LOOP task and filling out the questionnaires, participants were debriefed about the cover story and reimbursed for their participation. The whole procedure, including instructions, LOOP task, questionnaires and debriefing, took approximately one hour.

### Statistical analysis

#### Model-agnostic analysis

A model-agnostic analysis of participants’ performance expectations was conducted to illustrate basic task effects in our behavioral data. To this end, we calculated a repeated-measures ANOVA including the factors Trial (20 Trials), Ability condition (High Ability vs Low Ability), and Interference condition (Agent vs. No Agent).

#### Computational modeling

Reinforcement learning equations based on Rescorla-Wagner were used to capture trial-by-trial changes in performance expectation ratings (Rescorla & Wagner, 1972; Zhang et al., 2020). The most basic equation reads as follows (1):

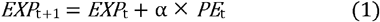

Here, “*EXP*_*t+1*_” stands for the expectation rating of the next trial, “*EXP*_*t*_” is the expectation rating of the current trial “*t*”, and “α” depicts the learning rate that functions as a weight on the prediction error “*PE*”. The prediction error “*PE*” is the difference between the received feedback “*FB*” and the current performance expectation “*EXP*_*t*_” (2):

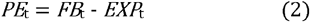

Note that the trial index “*t*” and trial-by-trial changes of expectation ratings are always understood within the respective experimental condition, since belief formation is specific for the estimation categories that are linked to the combinations of the levels of Interference × Ability. The basic equation was expanded in two ways based on previous studies (Czekalla et al., 2024; Müller-Pinzler et al., 2022; Schröder et al., 2024): First, different learning rates were introduced to give varying weight to prediction errors (PEs) based on their valence — differentiating between positive PEs (“better than expected”) and negative PEs (“worse than expected”). Additionally, a parameter *w* — a feedback weight factor — was introduced to lessen the impact of feedback near the scale’s limits. The latter was introduced based on the assumption that feedback that approaches the 0 or 100 percentiles may be less probable and therefore holds less informational value. This parameter *w* reduces the impact of PEs according to the feedback percentile. For this, it is multiplied by the relative probability density of the normal distribution “*ND*” of the possible feedback values, culminating in the extended equation (3) that serves as the base model for all models included in our model space:

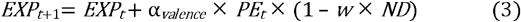

The final model space (see Figure 2B) contained five models. The first model (M1) assumed no influence of the Interference condition or attribution and only differentiated two learning rates for positive vs. negative PEs as shown in the equation above. All other models considered the Interference condition, but differed in their approach. A first set of two models did so by including the Inference condition, but without considering individual attributions per trial. The Interference Valence Model (M2) proposed four distinct learning rates based on PE valence and the level of Interference. The Interference Weighting Model (M3) assumed only two learning rates, separated by valence, but introduced an additional parameter *s*. This parameter also decreased the update, but only in trials of the Agent condition. The second and final set of two models assumed an influence of trial-by-trial attributions on updating, whether subjects believed an agent had manipulated the feedback. The Attribution Valence Model (M4) considered four learning rates based on PE valence and whether or not participants indicated that an agent had manipulated the feedback in the respective trial. The winning model, the Attribution Weighting Model (M5), proposed only two learning rates separated by PE valence and again considered a parameter *s*. In this model, the parameter *s* exclusively decreased learning rates for external attributions, so the model’s equations differed for trials with internal (4) vs. external attributions (5):

for internal attributions:

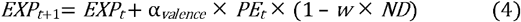

for external attributions:

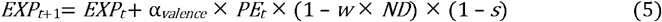

The model estimation resulted in largely uncorrelated estimates of learning rates and the parameters *w* and *s* (see Supplementary Table S3). For trials of the No Agent condition, where no attribution ratings were available, we assumed internal attributions throughout.

Based on previous studies that used the LOOP task, for this study, we deliberately did not consider simpler models that assume a singular learning rate across all conditions or different learning rates for each level of the factor Ability, since these model variants were regularly outperformed by models that differentiate learning rates for positive vs. negative PEs (Czekalla et al., 2024, 2021; Müller-Pinzler et al., 2022, 2019; Schröder et al., 2024).

#### Model fitting

For model fitting, the *rstan* package (Stan Development Team, 2021) for the software *R* was utilized. For all subjects, models were fitted individually with Markov Chain Monte Carlo (MCMC) sampling. Four MCMC chains were used and 3400 samples (thereof 1000 burn-in samples) were drawn, thinned by a factor of three. 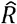 values for each parameter were inspected to check the convergence of the MCMC chains. Furthermore, the effective sample sizes (*n*_*eff*_) were verified to generally exceed 1500. This was done to check whether there were sufficient numbers of independent draws from the posterior distributions of the model’s parameters. After the successful model estimation, mean values per parameter and subject were calculated to summarize the posterior distributions.

#### Model selection

To determine which model best assessed the subjects’ updates of performance expectations, we estimated the pointwise out-of-sample accuracy for all fitted models within the model space, separately for each subject. For this, leave-one-out cross-validation (*LOO*; in this case, one trial per subject was left out) was utilized, and Pareto-smoothed importance sampling (*PSIS*) was applied by using the log-likelihood that was calculated from the posterior simulations of the estimated parameters (Vehtari et al., 2016). Sum *PSIS-LOO* scores for each model were used for model comparison (Supplementary Table S1). In addition, 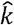 values were inspected - the estimated shape parameters of the generalized Pareto distribution. These values indicate the reliability of the *PSIS-LOO* estimates, and very few trials resulted in values that indicate unreliable scores (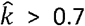, Supplementary Table S1). Further, Bayesian model selection (BMS) using *PSIS-LOO* values was conducted to identify the model that best described participant behavior at the sample level, considering the heterogeneity among participants. This provided the protected exceedance probability (*pxp*) for each model considered. The *pxp* is a metric that indicates the likelihood of a given model explaining the data in comparison to all other models in the model space. Additionally, it included the Bayesian omnibus risk (*BOR*), which serves as the posterior probability that the frequencies of all the considered models are equal (see Supplementary Figure S1 for BMS results).

#### Posterior predictive checks

To assess if the winning model captured the core effects in participant behavior, we repeated the model-agnostic analysis conducted on actual performance expectations with the data predicted by the winning model as a posterior predictive check. Specifically, we conducted a repeated-measures ANOVA on predicted performance expectations, with Trial, Ability, and Interference as within-subject factors.

#### Analysis of model parameters

Model parameters, that is, learning rates and attribution weight factors, of the winning model were analyzed on the group level using *R* Version 4.1.3. We compared learning rates for positive and negative PEs by calculating a Wilcoxon rank signed test. Attribution weight factors were compared against 0 using a one-sample *t*-test to demonstrate that attributions of feedback significantly influenced updates in performance expectations. Attribution weight factors of 0 indicate that causal attributions of the received feedback (internal vs. external) do not affect self-belief formation.

To investigate the associations of depressive symptoms and self-esteem with biases in learning and attributions, we calculated two valence bias scores for each subject. For the bias in learning, this was done by dividing the difference between learning rates for positive and negative PEs by their sum (Müller-Pinzler et al., 2022, 2019; Niv et al., 2012; Schröder et al., 2024):

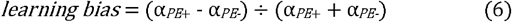

The bias in attributions was calculated analogously using the percentage of external attributions for positive and negative PEs instead of learning rates. To ensure comparability in the direction of the two biases (bias < 0: negativity bias, bias > 0: positivity/self-serving bias), the bias in attributions was multiplied by -1. Spearman correlations were then calculated between individual bias scores and PHQ-9 and SDQ-III sum scores, respectively.

## Acknowledgements

We want to thank Alicia Kroell for her invaluable assistance with data collection.

The research was funded by the German Research Foundation (Project-based funding for TK: KU 3955/3-1 and Temporary Positions for Principal Investigators LMP and SK: MU 4373/1-1; MU 4373/1-3; KR 3803/11-1; 3803/14-1) and the Medical Department of the University of Lübeck (J21-2018).

## Author contributions

A.V.M., A.S., D.S.S., N.C., S.K., and T.K. designed the research. A.S. and A.V.M. realized the experimental task and supervised data acquisition. A.V.M., A.S., and D.S.S. analyzed the data. A.V.M. and A.S. prepared the manuscript. All authors discussed the data analyses and interpretation of the results and reviewed and edited the manuscript.

## Data and code availability

The code and behavioral data for the statistical analyses are available at https://osf.io/2avsh/?view_only=5bd60291ceab4018b181eb26d98ca503.

