## Supplemental Material for "Self-esteem modulates beneficial causal attributions in the formation of novel self-beliefs"

**Contents**

**Supplementary Figures	2**

Figure S1: Bayesian Model Selection.	2

Figure S2: Simulated data demonstrating how the attribution weight factor of the winning model affects performance expectation trajectories.	3

**Supplementary Tables	4**

Table S1: PSIS-LOO scores.	4

Table S2: Posterior predictive check: repeated-measures ANOVA on simulated performance expectations	4

Table S3: Correlations of model parameters (winning computational model M5)	5

**Supplementary Notes	6**

Supplementary Note 1: Deviations from preregistration	6

**References	7**

### Supplementary Figures

#### Figure S1: Bayesian Model Selection.

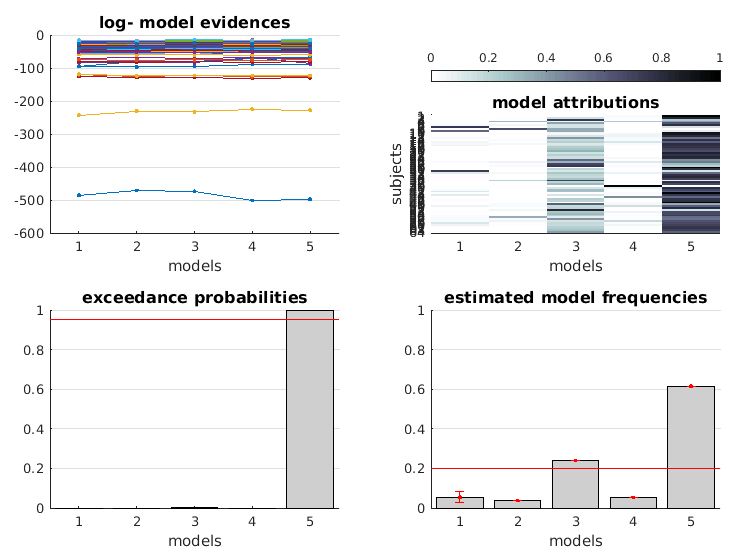

*Note.* Model 5 (Attribution Shrink Model) emerged as the winning model with a protected exceedance probability pxp = .999. and an estimated model frequency of 61.49.

#### Figure S2: Simulated data demonstrating how the attribution weight factor of the winning model affects performance expectation trajectories.

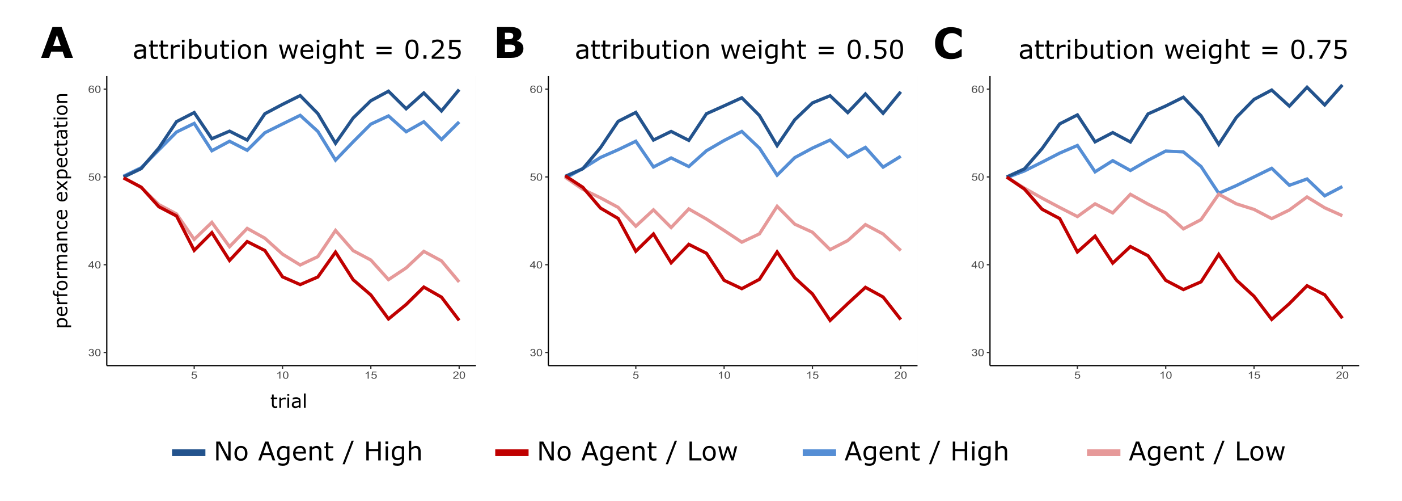

*Note.* These plots show how the attribution weight parameter in the winning model attenuates trial-by-trial learning in the Agent condition when feedback is attributed externally. It is important to note that this data was simulated based on the assumptions of an “optimal learner” in regard to this task. This means that expectations started at 50 in all conditions, the self-beliefs were updated according to the feedback, no counterfactual updates (e.g., a positive update following a negative prediction error) were considered, and only a small degree of noise was fitted onto the expectation ratings. The simulation assumed a learning rate for positive prediction errors of 0.15 and one for negative prediction errors of 0.2. A feedback weighting factor *w* of 0.9 was used. In the Agent condition, 40% of all trials were considered to be manipulated by the agent. **A** shows data with an attribution weight factor of 0.25, **B** of 0.5, and **C** of 0.75.

### Supplementary Tables

#### Table S1: PSIS-LOO scores.

| **Model** | **PSIS LOO** | **LOO SE** | **PSIS LOO Diff** | **LOO SE Diff** | **percent. *k̂* > 0.7** | **No. est. param.** |
| --- | --- | --- | --- | --- | --- | --- |
| Model 1 | -3366.86 | 525.62 | -122.55 | -8.12 | 0.76 | 7 |
| Model 2 | -3317.81 | 509.88 | -73.5 | -23.86 | 1.29 | 9 |
| Model 3 | -3268.48 | 515.33 | -24.17 | -18.41 | 0.92 | 8 |
| Model 4 | -3287.03 | 532.94 | -42.72 | -0.8 | 1.11 | 9 |
| Model 5 | -3244.31 | 533.74 | - | - | 0.92 | 8 |
| *Note.* PSIS-LOO = sum score of approximate leave-one-out cross-validation (LOO) using Pareto-smoothed importance sampling (PSIS); LOO-SE = standard error of PSIS-LOO; LOO-Diff = difference in expected predictive accuracy for all models in reference to the winning model with the highest PSIS-LOO (M5, Attribution Shrink Model; highlighted row) and standard error of these differences; percentage of *k̂* (estimated shape parameters of the generalized Pareto distribution) that exceed 0.7 (Vehtari et al. 2016); No. est. param. = number of parameters the model estimates (learning rates, initial expectation ratings for each condition, weight, and shrink factors where applicable). | | | | | | |

#### Table S2: Posterior predictive check: repeated-measures ANOVA on simulated performance expectations

| **Effect** | **F** | **df** | **p** | **generalized η2** |
| --- | --- | --- | --- | --- |
| Trial | 8.94 | 1, 63 | 0.004 | 0.02 |
| Interference | 1.34 | 1, 63 | 0.251 | 0.00 |
| Ability | 27.79 | 1, 63 | < .001 | 0.12 |
| Trial × Interference | 1.56 | 1, 63 | 0.216 | 0.00 |
| Trial × Ability | 62.80 | 1, 63 | < .001 | 0.02 |
| Interference × Ability | 10.92 | 1, 63 | 0.002 | 0.06 |
| Trial × Ability × Interference | 30.62 | 1, 63 | < .001 | 0.01 |
| *Note.* The pattern of results suggests that the winning model accurately captured the core effects in participant behavior. | | | | |

#### Table S3: Correlations of model parameters (winning computational model M5)

|  | **LR neg** | **attribution weight** | **feedback weight** |
| --- | --- | --- | --- |
| **LR pos** | *ρ* = -0.4,  *p_FDR_* = .005* | *ρ* = -0.06,  *p_FDR_* = .629 | *ρ* = 0.2,  *p_FDR_* = .216 |
| **LR neg** |  | *ρ* = 0.07,  *p_FDR_* = .629 | *ρ* = -0.25,  *p_FDR_* = .137 |
| **attribution weight** |  |  | *ρ* = -0.07,  *p_FDR_* = .629 |
| *Note.* *ρ* = Spearman’s rank correlation coefficient. Benjamini-Hochberg false discovery rate correction (FDR) was applied to correct *p*-values for multiple testing. The attribution weight and feedback weight parameters are not significantly correlated with the learning rates or with each other. | | | |

### Supplementary Notes

#### Supplementary Note 1: Deviations from preregistration

This study was preregistered on aspredicted.org (report #159493; https://aspredicted.org/n75w-npng.pdf). However, the methods used for analyzing the data diverged from the preregistered techniques in some aspects. This primarily relates to elements arising from the computational modeling process. The preregistered methods proposed a computational model that incorporates learning rates specifically designed to address valence-dependent updates in the Interference condition, aiming to explain participants' updating behavior through these learning rates (this corresponds to model M2; see Figure 2B and the methods section). Participants with higher PHQ-9 scores for depression were assumed to have lower learning rates in the positive Interference condition and higher learning rates in the negative Interference condition. This would indicate that participants with higher depression scores showed less self-serving updating. For the final publication, individual trial-wise attributions (whether the external agent’s interference was assumed or not) were incorporated into the winning computational model. This resulted in learning rates for updates following positive vs. negative prediction errors that were shared parameters between all conditions (compare Figure 1B). An additional model parameter, the attribution weighting factor *s*, was included to assign a weight to these learning rates whenever the participant assumed agent interference. As a result, learning behavior in these conditions could not be solely explained by learning rates as dependent variables. Alternatively, attribution weighting factors and the percentages of internal vs. external attributions following positive or negative prediction errors were analyzed. Further, self-esteem (SDQ-III; see methods section: questionnaires and debriefing) was included as an additional dependent variable to complement the score on depressive symptom severity. Instead of including self-esteem and depressive symptom severity as covariates, correlations with the scores for learning and attributional biases were calculated.
